# Sparse functional data analysis accounts for missing information in single-cell epigenomics

**DOI:** 10.1101/504365

**Authors:** Pedro Madrigal, Xiongtao Dai, Pantelis Z. Hadjipantelis

## Abstract

Single-cell epigenome assays produce sparsely sampled data, leading to coverage pooling across cells to increase resolution. Imputation of missing data using deep learning is available but requires intensive computation, and it has been applied only to DNA methylation obtained by single cell bisulfite sequencing. Here, sparsity in chromatin accessibility obtained by scNMT-seq is addressed using functional data analysis to fit sparsely sampled GpC coverage profiles of individual cells taking into account all the cells of the same cell-type or condition. For that, sparse functional principal component analysis (S-FPCA) is applied, and the principal components are used to estimate chromatin accessibility coverage in individual cells. This methodology can potentially be used with other single-cell assays with missing data such as scBS-seq, scNOME-seq, or scATAC-seq. The R package `fdapace` is available in CRAN, and R code used in this manuscript can be found at: http://github.com/pmb59/sparseSingleCell.

## Introduction

The cell is the basic biological unit of all living organisms. Recent developments allow genome-wide profiling of distinct regulatory layers at the single-cell level, some of them obtained in parallel (Chappell et al., 2018). For instance, single-cell nucleosome, methylation and transcription sequencing, or scNMT-seq, interrogates chromatin accessibility, methylation and RNA on individual cells (Clark et al., 2018). scNMT-seq and other multi-omics techniques such as scNOMe-seq have been used to profile the epigenome in mouse and human embryos during preimplantation development (Guo et al., 2017; Li et al., 2018), or in human cell lines GM12878 and K562 (Pott, 2017). After quality control and read alignment, single cells are normally clustered into different groups, and DNA methylation or chromatin accessibility in different regions of the genome (e.g., promoters, introns, enhancers, etc.) is compared between different cells or clusters. Challenges in single-cell epigenome data analysis include, but are not limited to: (1) Noisy data due to low amounts of starting material; (2) Poor coverage and sparse data (information across cells is normally pooled to increase resolution); and (3) Lack of computational approaches for data integration (Kelsey et al., 2017). Missing data is unavoidable even increasing sequencing to saturation (Clark et al., 2018).

In scNMT-seq, this means that data is obtained from different CpGs (methylation) and GpCs (chromatin) across different cells; i.e. data is sparsely distributed along the genome (Clark et al., 2018). Due to sparsity of the data we have many missing values at the single-cell level (Kelsey et al., 2017). In addition to cell pooling, two solutions have been proposed for DNA methylation (CpG coverage). The first uses a computationally intensive machine learning to predict missing values in single-cell methylation (Angermueller et al., 2017). The second approach uses a Bayesian hierarchical method to quantify methylation variation locally to cluster an impute missing data (Kapourani and Sanguinetti, 2018). None of these approaches have been used in single-cell chromatin accessibility.

As a proof of concept, here we apply sparse functional principal component analysis (S-FPCA) for the analysis of chromatin accessibility data obtained from scNMT-seq (Clark et al., 2018). Clustering is not approached here, and single cells should be clustered in advance using other packages such as Scasat (Baker et al., 2018) or scABC (Zamanighomi et al., 2018).

## Data

Data from 69 mouse embryonic stem cell (mESCs) was downloaded from GEO accession GSE109262. Processed GpC coverages were obtained from latest versions of Bismark, which includes NOME-seq functionality for scNMT-seq (Krueger and Andrews, 2011). We focus the analysis on chromatin accessibility GpCs instead on CpG methylation for two reasons. First, we get more GpC values than CpG for this experiment. Second, no tool has been proposed so far to model and reconstruct chromatin accessibility in scNMT-seq data.

## Methodology and Results

Functional Principal Component Analysis (FPCA) is well suited for data sparsely observed (Yao et al., 2005; Wang et al., 2015). This application was first inspired by the theoretical framework for sparse functional data analysis (S-FDA) described in (Kokoszka and Reimherr, 2017). It is important to note that smoothing is not applied to individual coverage profiles, but smoothing in S-FDA is based on pooling information from *N* cells. Imputed smooth coverages can be obtained only after information from all the cells (of same cluster) has been suitably combined. For each cell *n*, a nucleosome value in *C*_*n*_ GpCs can be observed. Then, we apply FPCA for sparsely or densely sampled random trajectories and time courses via the Principal Analysis by Conditional Estimation (PACE) algorithm included in the R package fdapace (Dai et al., 2017). Whether the data is considered sparsely or densely sampled is considered automatically by FPCA, the package’s main function for computing FPCA for dense or sparse functional data. We can examine the distribution of measurements along the domain of interest using a design plot (Yao et al., 2005) (**Fig. 1**). If they are notable gaps, we should use a suitably truncated domain for our analysis as to avoid potentially misleading boundary effects during smoothing.

**Fig. 1.**
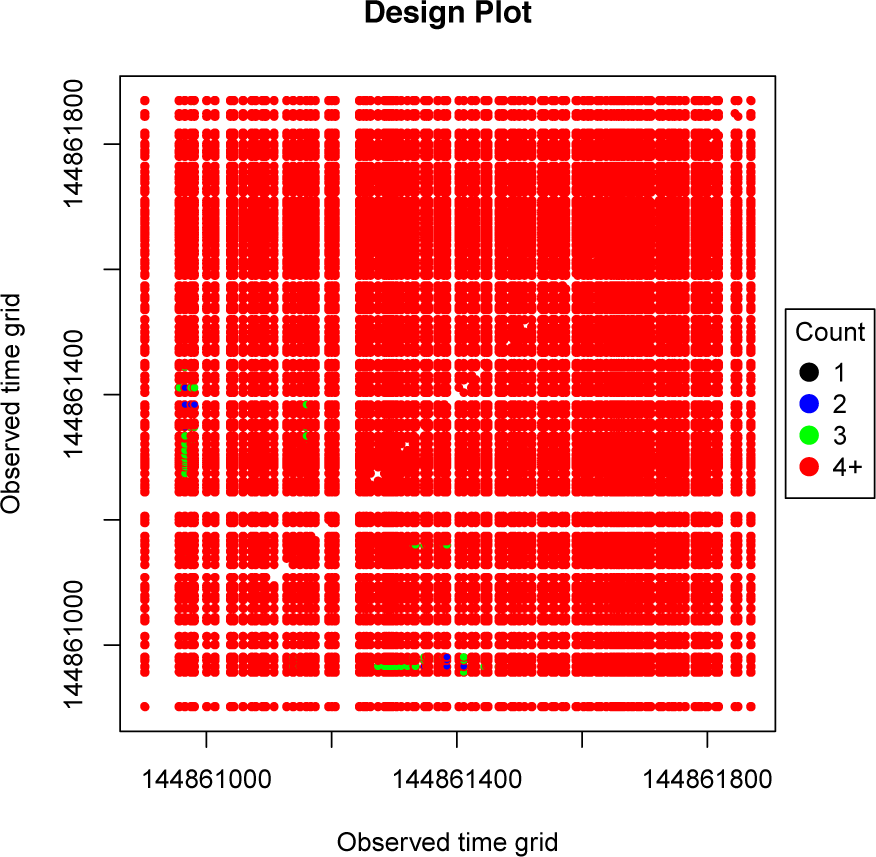
Design plot. The design plot showing the measurements times along the genome. The data show a relatively uniform spread and little to no clustering. The sparsity of the measurements before 144861000 suggest that using a left-truncated domain can help avoiding boundary effects. Diagonal measurements are not shown.

Assuming an unknown mean function *E{X*(*t*)*}* = *µ*(*t*) for the values of GpCs and an unknown auto-covariance function *cov*(*X*(*t*), *X*(*s*)) = *C*_*X*_ (*s, t*) where *s, t* are measurement points along the genome, the S-FPCA/PACE algorithm estimates the auto-covariance function *C*_*X*_ :

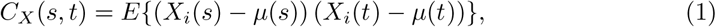

by pooling the raw subject-specific covariances together and smoothing the resulting point-cloud. Each subject-specific covariance is derived from the observed repeated GpCs measurements directly, whereas the within-sequence coverage measurements are allowed to be sparsely and irregularly made. The pooling then produces a point-cloud of raw covariances estimates between multiple different *s* and *t* measurements from each individual curve. The auto-covariance *C*_*X*_ is then estimated by estimating the smoothed surface. Given *C*_*X*_, its spectral decomposition can be written as:

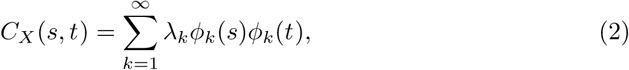

where the eigenfunctions *ϕ*_*k*_ are treated as the FPCA-generated continuous modes of variation - these are the Functional Principal Components (FPCs). The eigenvalues *λ*_*k*_ encapsulate the magnitude over variance of GpCs expression along the *k*-th principal component; they can also be used to determine the total percentage of variance along the *k*-th FPC (see scree-plot in **Fig. 2**). For any given *k* we can then use the corresponding eigenfunctions *ϕ*_*k*_(*t*) to compute *ξ*_*i,k*_, the projection scores associated with the *i*-th sample and the *k*-th component (Eq. 3) as:

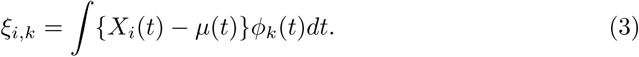

**Fig. 2.**
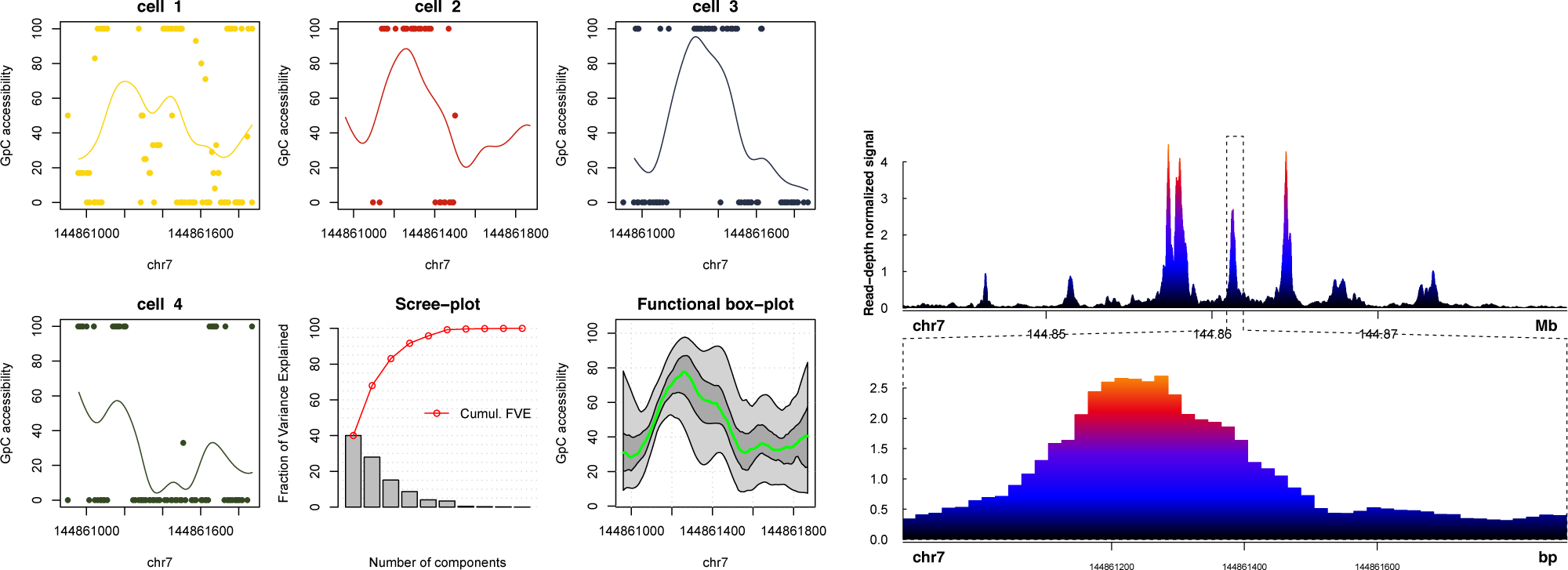
Sparse functional PCA applied to single-cell epigenomic data for 69 mouse ESCs. (Left) Visualization of predicted GpC coverage of chromatin accessibility for four cells based on the results from S-FPCA. The region analyzed corresponds to ±500 bp around the transcription start site of *Fgf4*. The scree-plot shows the proportion of variance explained by each functional principal component. The green line in the functional bag-plot (based on the function scores) corresponds to the functional median, the dark grey area to the area spanned by the curves within the 25th and 75-th percentile and the light grey to the area spanned by the curves within the 2.5th and 97.5-th percentile. **(Right)** Bulk DNase-seq data for *Mus musculus* C57BL/6 R1 cell line download from ENCODE (ENCFF974HWL). Zoomed region corresponds to the region used for sparse FPCA in scNMT-seq data. The visualization was generated with the R/Bioconductor package Sushi (Phanstiel et al., 2018).

Nevertheless, because of sparse and potentially irregular nature of the available data, direct numerical integration of Eq. 3 is potentially inaccurate or even unfeasible. We use a probabilistic approximation based on conditioning upon the observed data (Yao et al., 2005). The projection scores of the *i*-th sample and the corresponding *k*-th FPCs (Eq. 3) are then derived as:

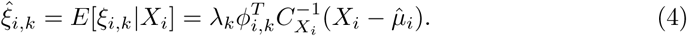

In that way, as *µ, C*_*x*_, *ϕ* and *λ* are produced using all the data available, the estimates for the scores *ξ*_*i,k*_ are optimal under Gaussian assumptions.

In summary, principal component scores *ξ* and FPCs *ϕ* are used to reconstruct individual trajectories, as well as create a functional box-plot. Importantly, for these operations, FPCA here does not rely on pre-smoothed coverages - e.g. as for ChIP-seq in previous applications of standard FPCA (Mateos et al., 2015; Madrigal and Krajewski, 2015).

To test the methodology we selected a region ±500 bp around *Fgf4*, a gene highly expressed in preimplantation epiblast of mouse embryos and in mouse embryonic stem cells. 58 out of 69 cells (84%) contained data values in these region. The pattern of measurement points can be seen in corresponding design plot (**Fig. 1**). The design plot shows that there is a uniform coverage across the genome’s base-pairs but also highlights that near the edges of the genome we have less measurements points. Because we want to avoid misleading boundary effect we partially truncate the start of the continuum examined by 6% (∼60 bp) in the subsequent analysis. Therefore, the corresponding FPCA estimates and descriptive statistics relate to this truncated domain. We run the function `FPCA` with parameters `‘maxK=30`, `nRegGrid=100`, `outPercent=c(0.06, 1)’`. Predicting GpC trajectories is possible in genomic regions of unobserved data (**Fig. 2**). The analysis in *Fgf4* shows also that the functional median of the 58 single cells resembles the signal from bulk DNase-seq performed in mESCs (**Fig. 2**).

## Conclusion

S-FPCA can be very useful to model and predict sparse single-cell epigenomes, as it uses information on all cells to estimate mathematical functions of epigenomic profiles containing missing data. Recovery of individual cell coverage could potentially help studying cellular heterogeneity, improve cluster accuracy, and facilitate integration of multi-omics data.

